# Comparison of QuPath and HALO platforms for analysis of the tumor microenvironment in prostate cancer

**DOI:** 10.1101/2025.05.16.654590

**Authors:** Wei Zhang, Qin Zhou, Jonathan V. Nguyen, Erika Egal, Qian Yang, Michael R. Freeman, Siwen Hu-Lieskovan, Gita Suneja, Anna Coghill, Beatrice S. Knudsen

## Abstract

QuPath, an open-source digital pathology platform, has gained widespread use for image analysis in biomedical research since its release in 2016. However, its reproducibility and reliability compared to commercial software, such as HALO, requires further validation, particularly for multiplex immunofluorescence (mIF) analysis. In this study, we performed a direct comparison of QuPath and HALO using a mIF-stained prostate cancer tissue microarray (TMA) inclusive of 192 unique cores. We evaluated performance across three key analytical modules: immune cell phenotyping, tumor infiltration with immune cells, and nearest neighbor analysis. Furthermore, we integrated QuPath with CytoMap, an open-source spatial analysis tool, to perform unsupervised clustering of immune cell infiltration—a feature not available in HALO. Our results demonstrated high concordance between two platforms, with correlation coefficients exceeding 0.89 for immune cell density, distance and pattern of cell organization in tumor microenvironment (TME). A neighborhood analysis using CytoMap was further performed and provided a more detailed spatial analysis of immune cell distribution across different prostate cancer grades. A significant increase of CD103+ T cell infiltration into TME was observed in prostate cancer, which might be associated with the expression level of its ligand (E-cadherin) in the tumor region. In conclusion, our findings validate QuPath as a robust and reproducible alternative to commercial platforms for fluorescence-based digital pathology. By demonstrating QuPath’s capability to perform high-quality quantitative analysis with additional flexibility for integration with external tools, our study underscores its potential for advancing tumor microenvironment research in translational oncology.

## Introduction

Prostate cancer, one of the most prevalent genitourinary malignancies, presents a significant public health challenge worldwide [1]. The disease originates in the epithelial cells of the prostate gland and can range from indolent and slow-growing to more aggressive forms with high metastatic potential [2]. Prostate cancer progression and prognosis are influenced by various factors, including genetic predispositions, environmental exposures, and, notably, the host’s immune system [3]. Immune cell infiltration within the tumor microenvironment is a critical component of the body’s defense against cancer. Infiltrating immune cells, including lymphocytes, macrophages, and dendritic cells, can either inhibit or promote tumor growth depending on their functional states and the signals they receive from the tumor and its microenvironment [4].

Determining the composition and activity of the immune cell infiltrate around solid tumors requires multiplexed antibody staining of tissues [5-7]. The quantification of immune cell phenotypes inside and surrounding cancer nests can be accomplished by image analysis. Multiple commercial analysis packages (e.g., HALO) are available in addition to the open-source QuPath platform [8-10]. However, it is uncertain whether commercial and open-source software will generate the same results, in particular for the quantification of rare immune cells.

HALO is a proprietary digital pathology image analysis platform developed by Indica Labs, which is ideal for automated, high-throughput, and user-friendly analysis, especially for well-defined workflows such as immune phenotyping, tumor infiltration, and spatial relationships. QuPath provides a free, highly customizable platform that supports integration with external tools like CytoMap, enabling unsupervised immune cell infiltration analysis, which is not available in HALO. However, it requires scripting and a manual setup of analysis pipelines. See detailed comparison between HALO and QuPath in Table 1.

**Table 1:** Feature comparison between HALO and QuPath.

| Feature | HALO (Indica Labs) | QuPath (Open-Source) |
| --- | --- | --- |
| Cost & Accessibility | Licensed, expensive | Free, open-source |
| User Interface | Intuitive GUI, minimal coding required | GUI with scripting support (Java/Groovy) |
| Image Formats | Supports various proprietary and open formats | Supports many formats, including Bio-Formats |
| Multiplexed Image analysis | Yes, built-in modules for immune phenotyping, tumor infiltration and spatial analysis | Yes, but requires manual setup and scripting for advanced analysis |
| Tumor infiltration analysis | Built-in tools for immune infiltration assessment | Requires custom scripting or integration with other tools (e.g., CytoMap) |
| Batch Processing | Yes, optimized for high-throughput workflows | Yes, but require scripting |
| Integration with other tools | Limited due to proprietary nature | Easily integrates with open-source tools (CytoMap, ImageJ, Python) |
| Customization & Flexibility | Limited customization | Highly flexible with scripting capabilities |
| Reproducibility | High but dependent on proprietary algorithms | High, fully transparent and modifiable workflows |

To assess the capabilities of open-source analysis platforms, we applied both the licensed HALO software (v3.6) and QuPath(v0.5.1) to analyze the same multiplexed TMA slides. We compared the output similarity across three analysis modules designated in HALO as “immune cell phenotyping”, “tumor infiltration”, and “nearest neighbor analysis”. Additionally, we integrated QuPath with another open-source platform, CytoMap, to explore phenotypic, unsupervised clustering of cells in the tumor microenvironment (TME), a feature not available in HALO. Moreover, single cell RNA sequencing analysis was utilized to validate the population of CD103+ T cell in prostate cancer samples.

## Materials and Methods

### Case collection

TMA slides of prostate cancers (192 cores) from 192 distinct patients were obtained from Moffitt Cancer Center in Tampa, FL. This TMA included 24 cores of Benign Prostatic Hyperplasia (BPH), 34 cores of Gleason Sum 6 (GS6, grade group 1), 42 cores of GS7 (3+4, grade group 2), 40 cores of GS7 (4+3, grade group 3) and 52 cores of GS8 (grade group 4), GS9 (grade group 5) and GS10 (grade group 5). One block of archival formalin-fixed and paraffin embedded high-grade prostate cancer was obtained from the Huntsman Cancer Institute (HCI) for investigation of CD103 and its ligand, E-Cadherin.

### Multiplexed immunofluorescence staining

TMA slides were immunostained at Moffitt Cancer Center using the Akoya Biosciences’ OPAL TM 7-Color Automation immunohistochemistry (IHC) kit (Marlborough, MA) on the BOND RX autostainer (Leica Biosystems, Vista, CA). At HCI, the tissue sections were stained with antibodies reactive with CD3 (Abcam, Catalog ID: AB16669), CD103 (Abcam, AB227697), E-cadherin (Abcam, AB40772) and pan-cytokeratin (Agilent, M3515) on the Leica Bond Rx autostainer (Leica Microsystems, Buffalo Grove, Illinois). The staining processes at Moffitt or Huntsman are same. Briefly, FFPE tissue sections were deparaffinized using the Bond dewax solution (AR9222) followed by sequential hydration steps in 100%, 95%, and 70% ethanol solutions. During antigen retrieval, tissue was immersed in ER1 -pH6 (AR9961-Bond TM Epitope Retrieval 1) for 30 minutes at 95°C. All subsequent steps were conducted using the automated OPAL multiplexed IF staining procedure, performed according to manufacturer protocols. This protocol consists of sequential staining and antibody removal cycles on the same tissue section until slides are stained with all antibodies from a panel. The OPAL 7-color kit uses tyramide signal amplification conjugated to individual fluorophores to detect targets (i.e., one fluorophore per immune marker). To evaluate the autofluorescence in the tissue, unstained slides were included as a negative control. After the antibody staining, slides were stained with 4′,6-diamidino-2-phenylindole (DAPI) to highlight the nuclei of cells in the tissue.

After staining was completed, TMA slides were imaged using the Vectra®3 Automated Quantitative Pathology Imaging System. Multispectral data were acquired from whole slide images at 20× magnification and at 20nm wavelength intervals between 420 and 690nm. Alternatively, HCI slides were scanned on the Zeiss Axioscan multispectral imaging platform with the appropriate filter cubes for each emission wavelength of individual OPAL fluorophores at 20×.

### Image processing by HALO and QuPath

Whole slide images of stained TMA slides were analyzed with the HALO Image Analysis Platform (Indica Labs, Albuquerque, NM) and QuPath. The first three steps including image loading, tissue segmentation, and tumor annotation are similar in both software programs. The DAPI and pan-cytokeratin (pCK) channels are used to obtain nuclear outlines and to separate epithelial from stromal regions. The separation of tumor and stroma was performed using a trained random forest supervised machine learning algorithm. Briefly, we manually annotated multiple regions of tumor and stroma in 5 TMA cores to train the RF model and applied it to all TMA cores to generate tumor outlines. Next, highplex module was used in HALO to detect nuclear outlines (**Supplementary Table 1**), classify cell phenotype, and identify cells in tumor or stroma. The threshold of positive versus negative stain separation was established for each marker based on visual assessment (**Supplementary Table 2**). A single manual threshold was established for each marker and applied to all the TMA cores to obtain the percentage of positive cells for a given marker/stain.

In QuPath, the dearrayer function was first applied to annotate each core with a unique label. Then we apply the STARDIST nuclear segmentation algorithm ^[11]^ to images at 20× magnification for detection of nuclei. To obtain cell borders, we expand the nuclear outlines by 5 pixels. The parameter settings of the STARDIST code for nuclear segmentation are available in Github (https://github.com/MarkZaidi/Universal-StarDist-for-QuPath/blob/main/Multimodal%20StarDist%20Segmentation.groovy). Single cell data including staining intensity of each channel were exported for further analysis. The same thresholds used in HALO for each marker were applied to single cell data generated from QuPath.

**Table 2:**
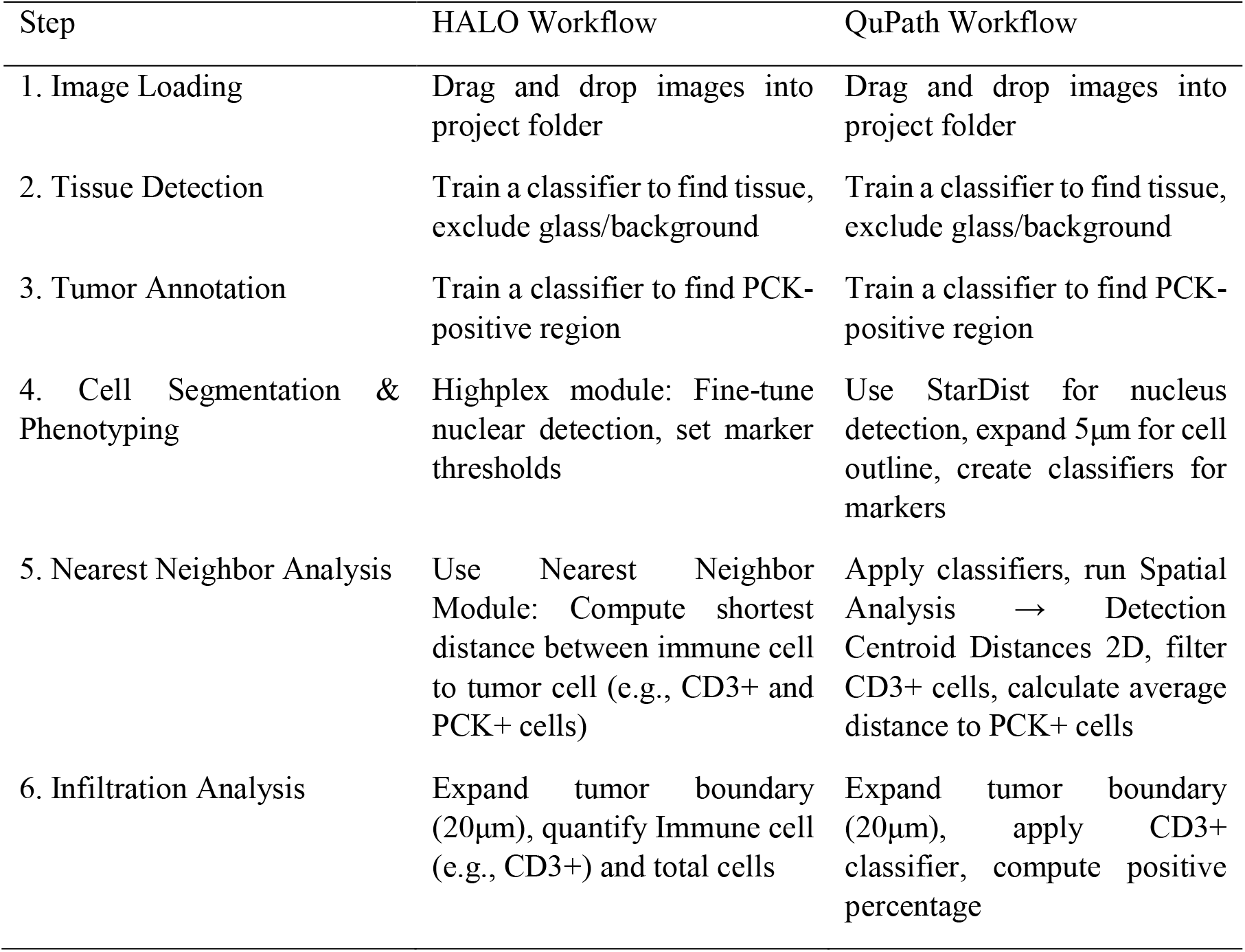
Comparison of HALO and QuPath Analysis Workflows.

### Spatial analysis

Nearest neighbor analysis. Using CD3+ cells as an example, we created separate CD3+ and pCK+ binary masks in HALO and performed the nearest neighbor analysis to obtain the average distance between CD3+ and pCK+ cells within each core. Similarly, in QuPath, we generated cell masks and applied both CD3+ and pCK+ classifiers to all cells. We then conducted spatial analysis using Detection Centroid Distances 2D to determine the shortest distance between CD3+ and pCK+ cells.

Tumor Infiltration module. To analyze the infiltration of immune cells into the tumor regions, we expanded the tumor outline by 20 µm and determined the number of positive cells for each marker. This was achieved either by applying a positive cell filter in HALO or a binary classifier in QuPath to count cells with the expanded tumor region. Based on nuclear segmentation in the DAPI channel, we can obtain both positive and total cell counts.

Neighborhood analysis. We use CytoMap, a MATLAB-based histo-cytometric multidimensional analysis pipeline designed for spatial analysis of segmented cell objects [12] to analyze the output from the QuPath. Using the coordinates of each cell of interest, CytoMAP employs diverse statistical approaches to extract and quantify information related to the positions of cells, cell-cell associations, and overall tissue structure. By grouping cells into local neighborhoods, CytoMap reveals patterns of cellular composition and tissue structures [12]. We imported single-cell data of immunofluorescent staining features with XY coordinates into CytoMAP and performed an unsupervised cluster analysis into three groups using the Self-Organizing Map method. We displayed the three groups of cell populations as tissue regions using CytoMap and determined the relationship of the tissue regions to the tumor mask. Next, we performed a cell neighborhood analysis within each region after classifying the regions as tumor, tumor-adjacent stroma, and tumor-distant stroma. The percentage of immune cells within each region was compared across different Gleason scores in prostate cancer cases.

### Single cell transcriptome data analysis

Three single-cell RNA-seq datasets from published studies were integrated [13-15]. Before integration, doublets and triplets were removed using DoubletFinder function in Seurat R package in each dataset [16]. In each dataset, 2000 highly variable genes were selected, and subjected to Principal Component Analysis (PCA). Cells with < 500 genes or with a mitochondrial gene expression content > 20% were removed (nFeature_RNA > 500 & percent.mi < 20%). Then, a total of 36419 primary prostate cancer and 23607 castration-resistant prostate cancer cells were further analyzed. Canonical correlation analysis was applied to adjust systemic biases from individual datasets (Seurat R package (version 4.3.0)) [17]. ComBAt function in SVA R package was used to remove batch effects during integration [18]. Then, we performed Uniform Manifold Approximation and Projection (UMAP) analysis to identify cell sub-populations [19]. Clustering was performed for integrated expression values using FindNeighbors function in Seurat package with the resolution of 0.8. The following known markers were used for cell type annotation: Epithelial/Tumor: EPCAM, KRT8; T cells: CD3E, CD3D, TRBC1/2, TRAC; B cells: CD79A/B, JCHAIN, IGKC, IGHG3; Myeloid cells: LYZ, CD86, CD68, FCGR3A; Endothelial cells: CLDN5, FLT1, CDH1, RAMP2; Fibroblasts: DCN, C1R, COL1A1, ACTA2; Mast cells: TPSAB1. Seurat R package was applied to show the expression of CD103 in the integrated set of single-cell RNA-Seq data.

### Statistical analysis

Scatter plot and linear fitting methods were used to compare the correlation between HALO and QuPath outputs. Box plots plots were utilized to visualize differences between Gleason Sum groups. In each box plot, the mean value is represented by a black line, and the whiskers extend to the 25th and 75th percentiles of the data distribution. All visualizations were generated using OriginLab Pro 2024b. For comparisons between two groups, t-tests were conducted, with a p-value threshold of 0.05 set to determine significance. Additionally, Pearson correlation analysis was employed to evaluate data correlations across statistical software outputs in each prostate core.

## Results

### Summary of Image analysis workflow

TMA slides were stained with panel 1 (CD3, CD8, PDL1, PD1, and FoxP3) and panel 2 (CD20, CD103, Delta-TCR, TIM3, and CD164) to identify B-cell and T-cell subsets. Images were first analyzed with the commercial HALO image analysis software. This software requires that the user provides parameters for nuclear segmentation, cell-wise measurements of average pixel intensity for each fluorophore, and the threshold for separation of positive and negative cells. Single cell staining data were collected in tumor and stromal regions from each TMA core using the pCK mask to map the tumor glands (i.e., to identify individual cells). The single cell data were analyzed with the HALO “nearest neighbor” and “tumor infiltration” modules. Next, we determined whether the open-source QuPath software, provides the same results as the HALO software package. We applied QuPath to the same images from the Vectra®3 slide scanner and followed an identical workflow to the one for data generation with HALO, except for the cell detection algorithm. A major difference between the two platforms is the pre-trained deep-learning network used for nuclear segmentation. Therefore, we compared the numbers of cancer cells detected by HALO or QuPath in each core and found a high correlation between the two platforms (SF 1).

We also compared results of other metrics between HALO and QuPath (**Fig.1**), using the HALO modules for immune cell phenotyping, nearest neighbor analysis (i.e. the shortest distance between immune and tumor cells), and tumor infiltration (i.e. the immune cell percentage in tumor adjacent regions). The detailed workflow is illustrated in **Fig. 1**.

**Figure 1.**
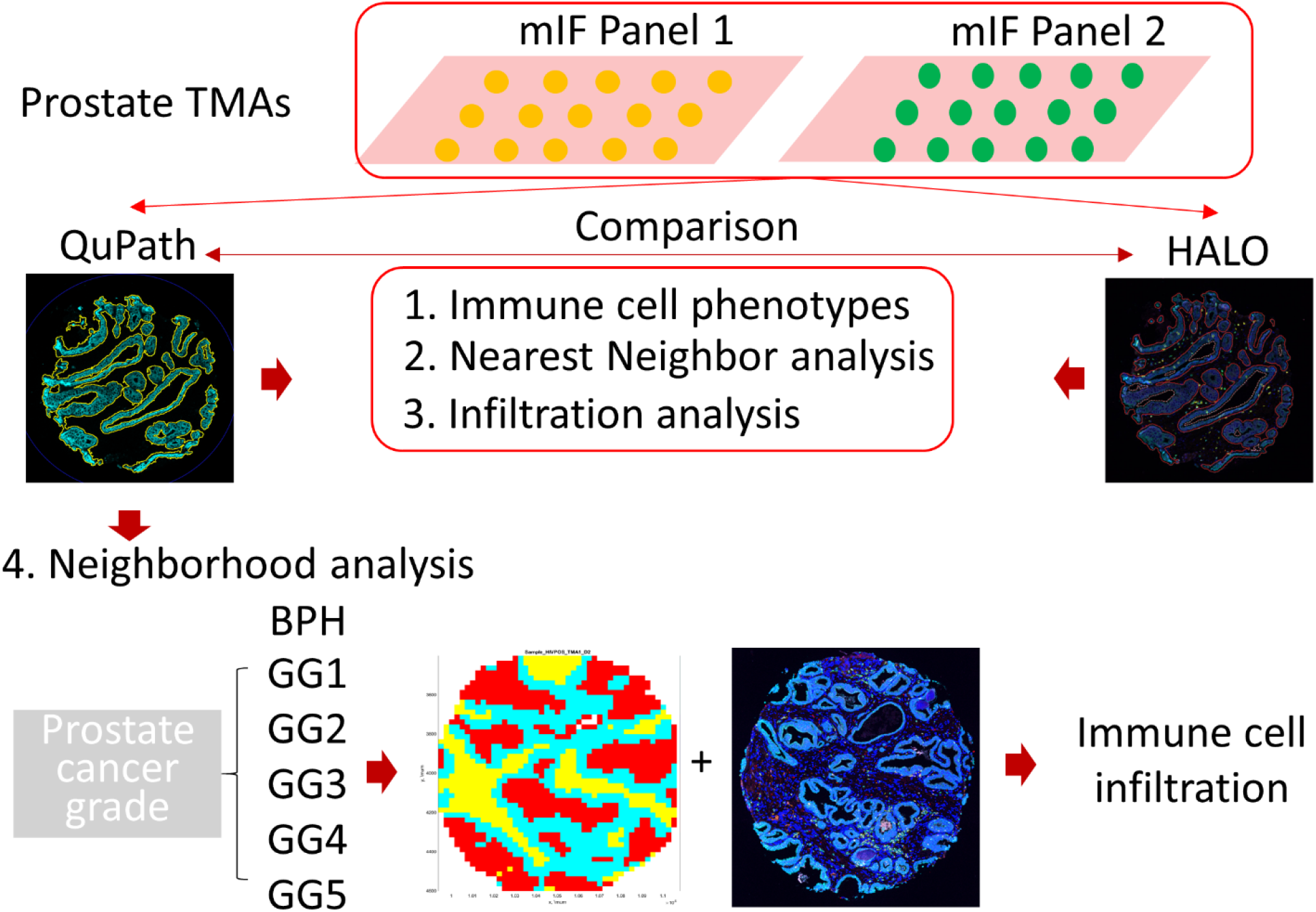
Workflow for comparison of QuPath and HALO image analysis results. Prostate cancer TMAs were stained with antibody panel 1 (CD3, CD8, PD-1, PD-L1, FoxP3, pCK), or with antibody panel 2 (CD20, CD103, Delta-TCR, TIM3, CD164, pCK). Scanned mIF images were analyzed using either QuPath or HALO image analysis software packages. The processing of images encompasses multiple steps, including tissue segmentation, cell detection, and quantification of immune cell populations. Quantitative measures of tumor regions and immune cell percentages can be obtained through analysis with HALO or QuPath, but unsupervised neighborhood analysis is only possible using a combination of QuPath and CytoMap.

### Comparison of immune cell phenotyping in HALO and QuPath

The tumor area masked by pCK staining was measured by HALO or QuPath in all TMA cores, revealing a strong linear correlation of 0.91. This indicates a high level of agreement between QuPath and HALO in pixel-level segmentation (**Supplementary Fig. 1**). Next, we calculated the percentages of positive immune cells for each marker by distinguishing positive and negative cells based on a manually defined threshold of signal intensity (**Supplementary Table 2**). Percentages of positive cells were calculated using either the number of cells inside the cancer mask or the total number of cells in the core as denominators. Pearson correlation analysis showed strong concordance (r > 0.89) for markers in panel 1 (CD3, CD8, PD-1, PD-L1, FoxP3, pCK) (**Fig. 2B**). Correlation coefficients for panel 2 markers (CD164, TIM3, and Delta-TCR) were slightly lower, likely due to their relatively low positive cell percentages (**Fig. 2D**). Similarly, correlation coefficients for pCK positive regions were slightly reduced (> 0.8) but still demonstrated good agreement (**Supplementary Fig. 2**). These findings confirm the reliability of QuPath-generated data and support its suitability for comprehensive immune cell population analysis.

**Figure 2.**
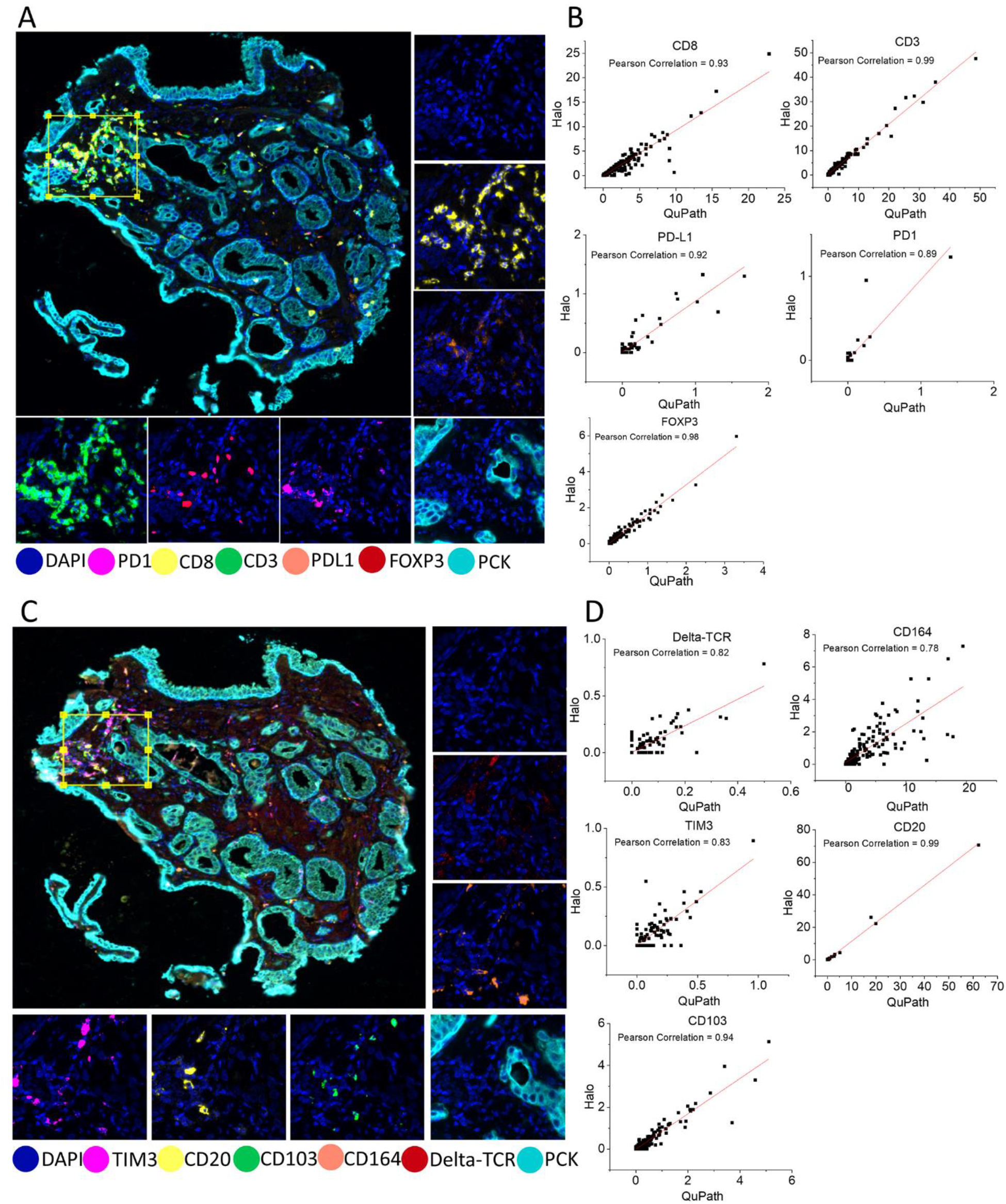
Comparison of immune cell phenotypes measured by HALO or QuPath. **A**. Panel 1, composed of antibodies reactive with PD1, CD8, CD3, PDL1, FoxP3, and pCK. **B**. Pearson correlation analysis between QuPath and HALO using the same signal intensity thresholds in HALO and QuPath to distinguish positive and negative pixels for analysis. **C**. Panel 2, encompassing markers TIM3, CD20, CD103, CD164, Delta-TCR, and pCK. **D**. Pearson correlation analysis between QuPath and HALO using the same signal intensity thresholds in HALO and QuPath to distinguish positive and negative pixels for analysis.

### Comparison of spatial analysis in HALO and QuPath

First, we compared the results from the tumor infiltration analysis between the two platforms. Using a concentric 20µm expansion of the tumor mask, we calculated immune cell density within the expanded region (**Fig. 3A**). We focused on the three most abundant immune cell markers—CD3, CD8, and CD103. Pearson correlation analysis revealed a strong agreement for CD3 and CD8 cells, with CD103 showing a slightly lower correlation coefficient of 0.89 between the software programs (**Fig. 3B**).

**Figure 3.**
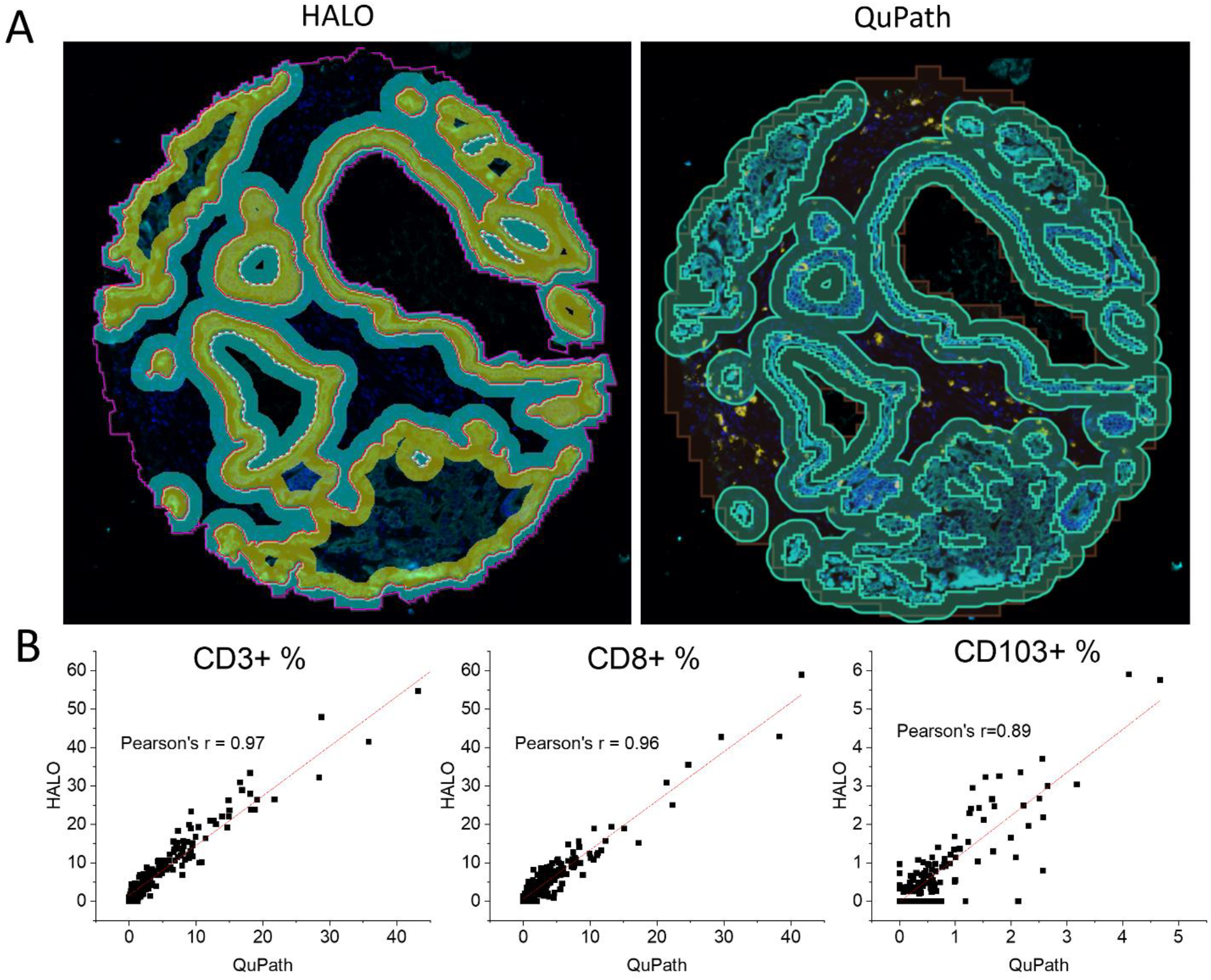
Comparison of infiltration analysis by HALO and QuPath. A. Tumor expansion of 20µm in HALO and QuPath. B. Linear fitting of scattering plot between HALO and QuPath output

In HALO, nearest neighbor analysis computes the shortest distance between the centroids of the two cell nuclei, for example a nucleus of a CD3+ cell and another of a pCK+ cell. This distance can also be measured in QuPath. Additionally, QuPath allows for the calculation of the shortest distance between the centroid and the outline of the tumor mask (cell-to-mask distance). **Fig. 4A** illustrates an example of results from nearest neighbor analysis with HALO. HALO automatically calculates the average distance and does not allow to retrieve individual distance measures. In contrast, QuPath allows to construct a histogram of pairwise distances (**Supplementary Fig. 3)**. Our analysis (**Fig. 4B**) demonstrates a high concordance for nearest cell-to-cell distance measurements (correlation > 0.90). As expected, the cell-to-mask distance is slightly less compared to cell-to-cell distance in QuPath for all three markers (**Supplementary Fig. 4**).

**Figure 4.**
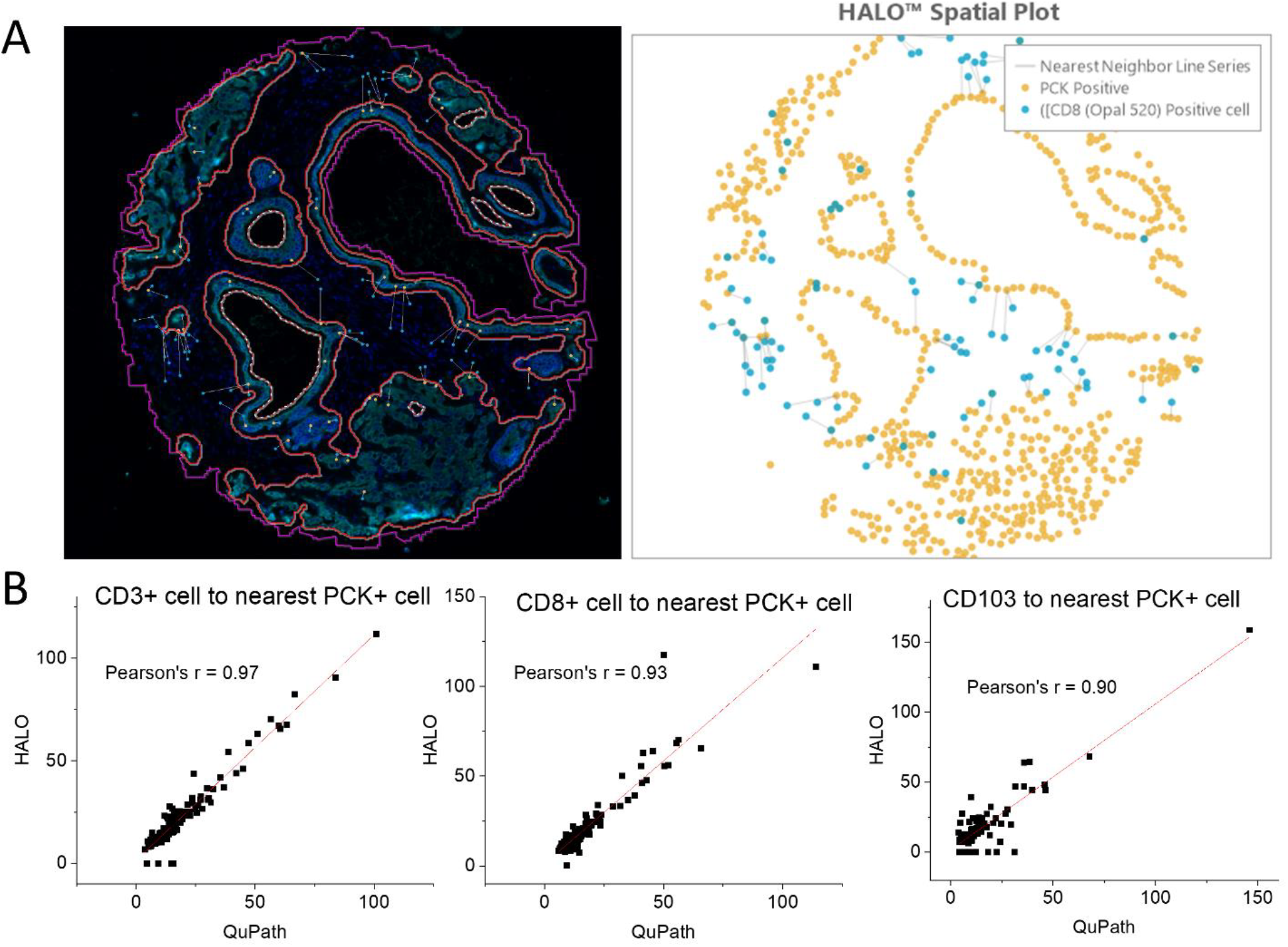
Comparison of nearest neighbor analysis by HALO and QuPath. A. Illustration of nearest neighbor analysis in HALO. B. Linear fitting of scatter plot between HALO and QuPath outputs.

### Neighborhood analysis in Cytomap

Next, we analyzed immune cell distributions in benign prostate and prostate cancer tissues across different Gleason grades. Using CytoMap, we performed a neighborhood analysis by unsupervised clustering of single-cell data into three regions (R1, R2, and R3). The percentage of pCK+ cells in each region revealed that R1 contains the largest number of benign or cancerous epithelial cells, R2 comprises the area adjacent to tumor and R3 is far away from the tumor (**Fig 5A**). Therefore, we named R1 as epithelium/cancer region, R2 as adjacent stroma and R3 as distant stroma. The relative percentage of immune cells within each region was calculated for comparison. We observed an increased presence of immune cells in the tumor-adjacent stroma of prostate cancer compared to benign prostatic hyperplasia (BPH) (**Fig 5B-C**). Notably, CD103+ cell infiltration increased significantly from 32% in BPH to over 52% in prostate cancer samples (p<0.05) (**Fig. 5C**). However, the proportion of CD103+ cells among different Gleason groups is not significant.

**Figure 5.**
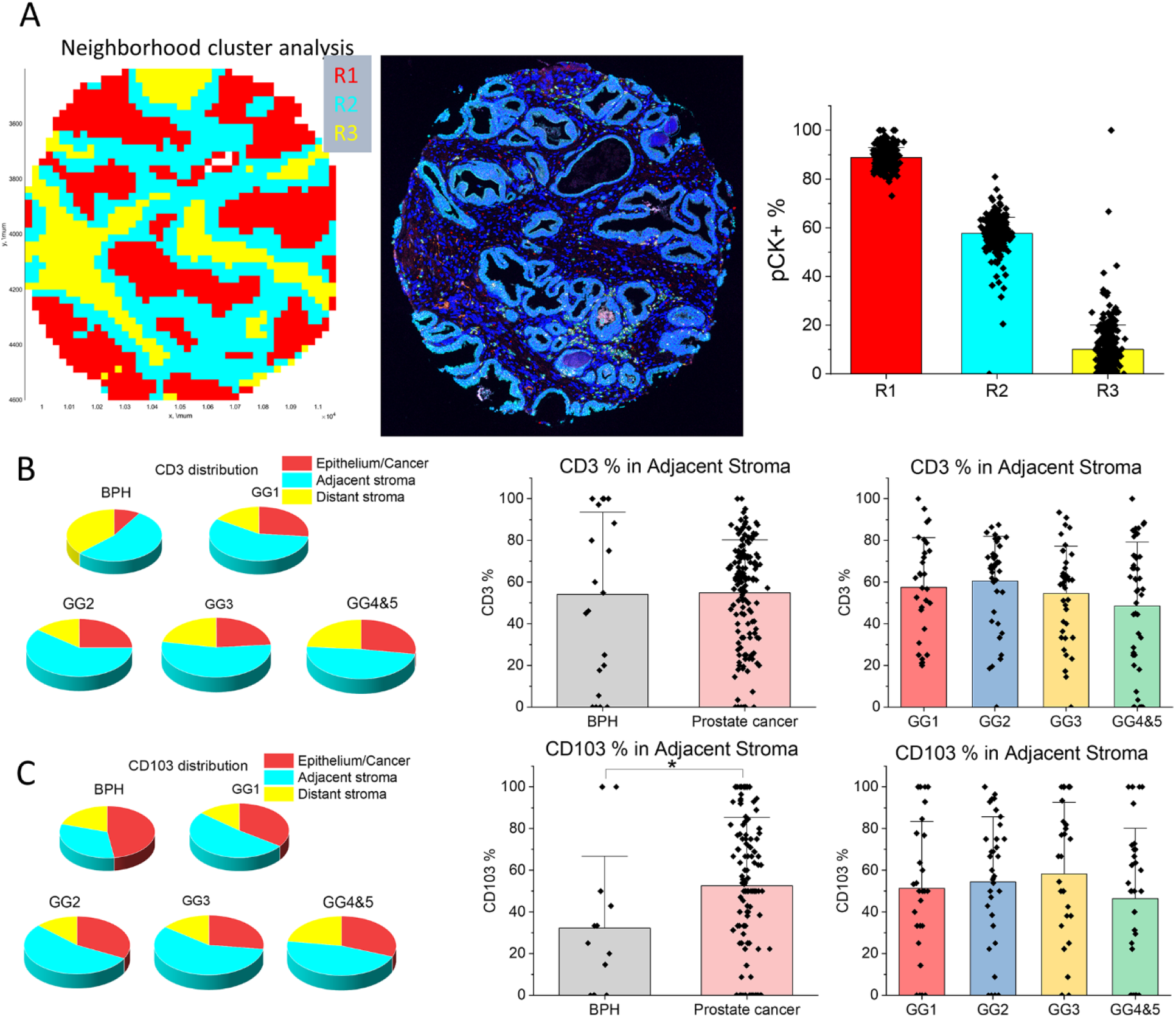
Comparison of immune cell densities. A. Example of neighborhood cluster analysis, wherein the cells in the entire core are clustered using an unsupervised approach into three groups based on antibody staining and cell-to-cell distance parameters. Visualization of the location of cells in the clusters generates three regions, a region of benign epithelium in BPH cores or a cancer region in cancer cores, a region adjacent to cancer and a region distant from the cancer. B. Relative distribution of CD3+ cells in three regions and comparison of CD3 percentages in adjacent stroma between BPH and prostate cancer or among different Gleason grade cases. C. Relative distribution of CD103+ cells in three regions and comparison of % CD103 in adjacent stroma between BPH and prostate cancer or among different Gleason grade cases. Significant differences (p < 0.05) are indicated by a *.

### Characterization of CD103 positive cells

Our study reveals a difference of CD103+ cells in the intra-tumor cell density and the spatial analysis between BPH and prostate cancer patients, prompting us to further characterize CD103+ cells in prostate cancer. We analyzed existing, single-cell RNA sequencing data from three public prostate cancer datasets (GSE141445, GSE157703 and GSE137829). A UMAP plot of *CD103+* gene expression at the RNA sequencing level (**Supplementary Fig. 5**) shows that the majority of *CD103+* cells express *CD3D* and *CD3E* (T cells markers) or *EpCAM, CK8, CDH1* (epithelial markers). CD103 is a ligand for E-cadherin, which is expressed on the surface of benign and most malignant prostate epithelial cells [20]. Since CD103+ immune cells are significantly increased in prostate cancers, we aimed to determine whether E-cadherin expression attracts CD103+ T cells. To investigate this question, we selected a high-grade prostate cancer case from HCI with cancer regions displaying low- and high E-cadherin levels (**Fig. 6A**). The case was stained it with CD103, CD3, E-cadherin, and pCK. We observed that the expression level of E-cadherin varied across cancer regions. We selected five regions of interest (ROIs) with high E-cadherin expression and five ROIs with low E-cadherin expression to compare the presence of CD103+ T cells (**Fig. 6B**). In this prostate cancer case, the percentages of CD3+ and CD103+ immune cells were significantly higher in areas with elevated E-cadherin levels. This suggests a possible role of the E-cadherin in the attraction of CD103+ T cells.

**Figure 6.**
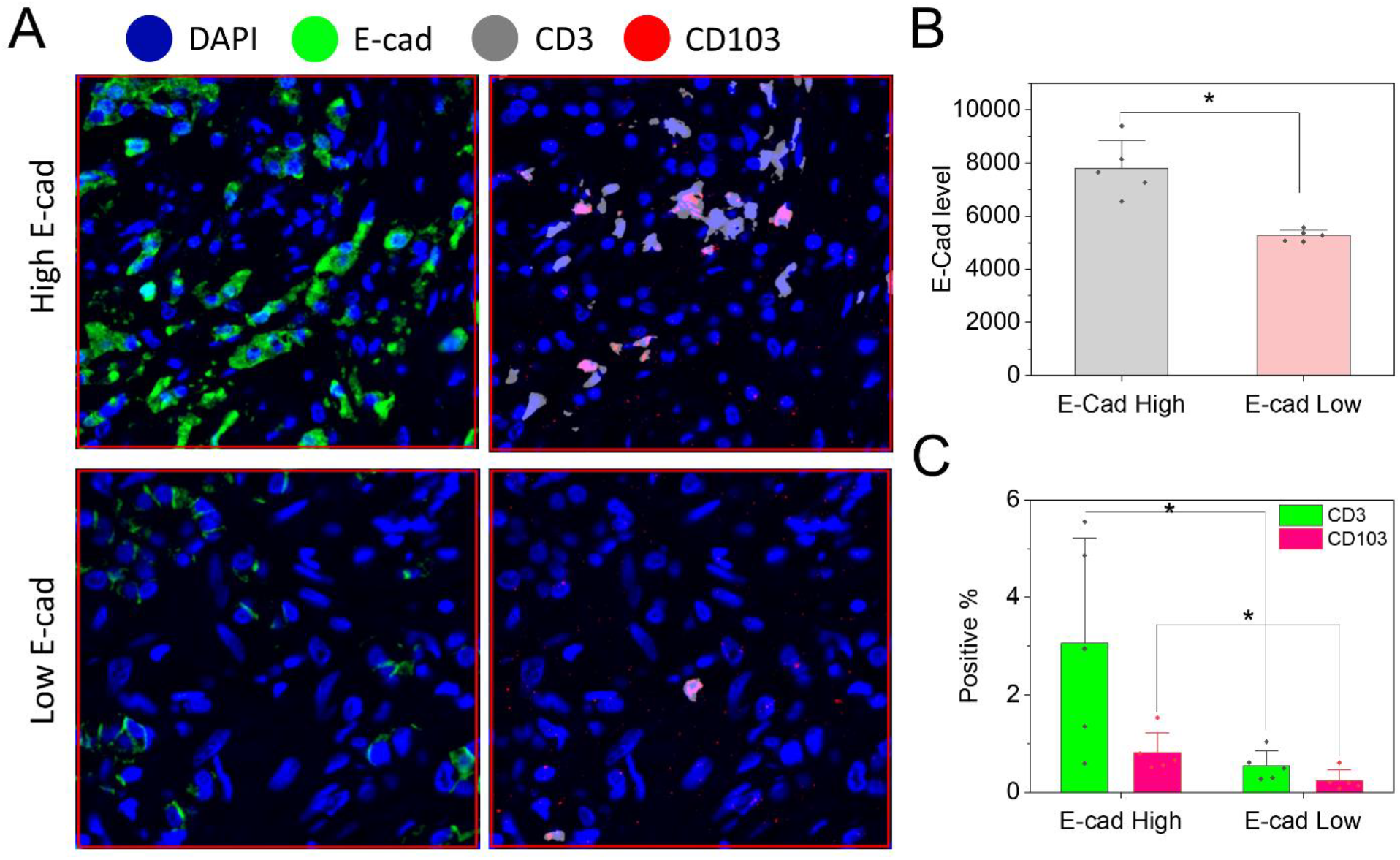
Frequencies of CD103+ cells in the proximity of prostate cancer cells with high-versus low E-Cadherin expression. One prostate cancer slide from a radical prostatectomy with regions of Gleason grade 4 + 5 tumor was stained with a panel of CD103/CD3/E-cadherin/pCK. A. Comparison of CD103+ cells in high-versus low E-cadherin expressing cancer regions. Cancer cells were identified based on pCK expression. **B**. E-cadherin expression levels in 5 ROIs from E-cadherin-high and E-cadherin low tumor regions. **C**. Percentage of CD3 and CD103 relative to tumor cells in E-cadherin-high versus E-cadherin-low tumor regions.

## Discussion

To our knowledge, our study provides the first report of a direct comparison between tumor immune staining output generated by the commercial HALO software package versus the open-source QuPath software. Three modules we compared all demonstrated very high agreement, which validates the use of open-source software QuPath to obtain accurate data from immunofluorescent tissue images. Moreover, the flexibility of the open-source platform allowed us to integrate with other open-source software tools for deeper analysis, such as CytoMap [21]. The combined QuPath – CytoMap functionality permitted an unsupervised neighborhood analysis, separating tumor regions for analysis of immune cell infiltrations. In addition, we observed the association of CD103+ T cells with E-cadherin positive prostate cancer cells.

QuPath has been widely used in digital pathology analysis since its initial release in 2016. Its reproducibility compared to commercial platforms has been extensively evaluated, particularly in bright-field imaging. For example, Balazs et al. assessed Ki67 expression using HALO, QuantCenter (3DHistech), and QuPath, demonstrating excellent reproducibility both within and across digital image analysis platforms [22]. Another study compared Definiens Tissue Studio®, inForm®, and QuPath for the analysis of clinical breast cancer markers (ER, PR, HER2, EGFR, and CK5/6) in tissue microarrays, finding that QuPath exhibited the highest correlation and agreement with manual scoring [23]. Our study further highlights the robustness of QuPath by demonstrating its high reliability for fluorescent image analysis, reinforcing its utility across both bright-field and fluorescence-based imaging applications.

Spatial technologies, such as spatial proteomics and transcriptomics, can reveal novel patterns of immune cell organization that may be linked to prognosis and response to treatments [24]. A study used Slide-seqV2, a spatial transcriptomic technique, to preserve tissue architecture and cell-to-cell proximity relationships, and confirmed the colocalization of a CD16-negative NK cell signature and CCL5 transcript expression in prostate cancer [25]. QuPath can be used as an open-source alternative to examine the spatial-temporal dynamics of immune cell types infiltrating tumors. For example, one study found that the density of tumor-associated macrophages (TAMs) decreased over the course of tumor growth and in response to treatments [26]. Our neighborhood analysis shows that significant difference of CD103+ cells in BPH versus prostate cancer, which prompted us to further characterize these cells. Several studies claimed that CD103+ immune cells are a subtype of T cells [27, 28], however other studies demonstrated CD103 expression in myeloid cells [29]. The abundance of CD103+ cells is associated with favorable prognosis and survival in different cancer types, including colorectal cancer [30], glioblastoma [31] and prostate cancer [32]. In prostate cancer, CD103+ immune cells are also predictive of improved response to treatment with androgen-ablative therapy [32]. Our analysis confirmed a high co-localization between CD103+ and CD3+ cells (**Fig. 6A**), conforming that CD103+ cells in our prostate cancer cases are indeed a subset of T cells. This is supported by single-cell RNA sequencing data, which revealed that a large percentage of CD103+ cells express the T cell marker, CD3. However, expression of CD103 mRNA is also observed in a population of epithelial cells. Interesting, CD103+ epithelial cells are not detected in the prostate tissue from our study, suggesting differences in the regulation of CD103 protein synthesis or turnover in immune and epithelial cells.

A recent study revealed that the binding of CD103 on tumor-infiltrating lymphocytes to immobilized recombinant E-cadherin induces the polarization of cytolytic granules, whereas degranulation requires the co-engagement of CD103 and the T-cell receptor. Thus, CD103 possesses a unique costimulatory role in tumor-specific cytotoxic T-cell activation by providing signals for T-cell effector functions to target and lyse cancer cells [33]. Another study revealed that the loss of E-cadherin inhibits the antitumor activity of CD103+ immune cells in melanoma. This study encourages drug development to increase the activation of CD103+ immune cells in E-cadherin expressing solid tumors [34]. Our findings suggest that higher levels of E-cadherin protein expression are associated with increases of CD3+ and CD103+ immune cells in the tumor region. While this implies an opportunity for activation of CD103+ T cells by E-cadherin, the CD103+ cells are distant from cancer glands and would require to be stimulated for tumor infiltration to exert their ant-tumor activity. Our data also support a potential scenario in which the activation of CD103 occurs by benign prostate cancer glands in the tumor region that express high E-cadherin levels with subsequent tumor infiltration of the CD103+ activated immune cells. The variability in E-cadherin expression across different regions of the tissue highlights the complexity of cell-cell interactions and their impact on immune cell localization and function.

In conclusion, this study provides the first direct comparison of HALO and QuPath for multiplex immunofluorescence image analysis, demonstrating that QuPath achieves highly reproducible results comparable to the commercial HALO software. The three analytical modules assessed—immune cell phenotyping, tumor infiltration, and nearest neighbor analysis—all showed strong agreement between the two platforms, supporting the reliability of QuPath for fluorescent tissue image analysis. Additionally, we leveraged the flexibility of QuPath by integrating it with CytoMap, enabling unsupervised immune infiltration analysis beyond HALO’s capabilities. Our study also highlights the role of CD103+ cells in the prostate cancer microenvironment, suggesting a potential association with E-cadherin expression. These findings underscore the value of open-source platforms for spatial immune profiling and their potential application in translational cancer research. Future studies incorporating larger datasets and spatial transcriptomic technologies could further enhance our understanding of immune cell interactions in prostate cancer.

## Authorship

BK and WZ designed the concept, WZ, QZ, JVN and EE performed and collected the data, QY and MRF provided the statistical analysis, BSK and WZ wrote the manuscript, GS, AC, SH and BSK revised the manuscript, all authors read and approved the final version of the manuscript.

## Ethics statements

The study involving humans were approved by Moffitt Cancer Center Institutional Review Board University of Utah Institutional Review Board. The studies were conducted in accordance with the local legislation and institutional requirements.

## Acknowledgements

This work was partially supported by the National Cancer Institute (UM1CA181255). We acknowledge the generous support by the Department of Pathology and the Huntsman Cancer Institute. We thank Jeff Stanley and the Biospecimen and Molecular Pathology Shared Resource for fluorescence staining. We acknowledge the direct financial support for the research reported in this publication provided by the Huntsman Cancer Foundation and the Experimental Therapeutics Program at Huntsman Cancer Institute. Research reported in this publication utilized the Biorepository and Molecular Pathology Shared Resource at Huntsman Cancer Institute at the University of Utah and was supported by the National Cancer Institute of the National Institutes of Health under Award Number P30CA042014. The content is solely the responsibility of the authors and does not necessarily represent the official views of the NIH.

