## Supplemental material for "Comparison of QuPath and HALO platforms for analysis of the tumor microenvironment in prostate cancer"

**Supplementary Table 1**. Nuclear detection parameters in HALO

| Parameter | Setting |
|---|---|
| Nuclear Contrast Threshold | 0.508 |
| Minimum Nuclear Intensity | 0.095 |
| Maximum Image Brightness | 1 |
| Nuclear Segmentation Aggressiveness | 0.523 |
| Fill Nuclear Holes | False |
| Nuclear Size | 11.3, 571.7 |
| Minimum Nuclear Roundness | 0 |
| Number of Nuclear Dyes | 1 |
| Nuclear Dye 1 | DAPI |
| Nuclear Dye 1 Weight | 1 |

**Supplementary Table 2.** Thresholds for T cell and Myeloid panel.

| Marker | Thresholding | Compartment |
|---|---|---|
| Delta TCR Opal 520 | 1 | Cytoplasm |
| CD20 Opal 620 | 6 | Cytoplasm |
| CD103 Opal 650 | 3 | Cytoplasm |
| CD164 Opal 540 | 3 | Cytoplasm |
| TIM3 Opal 570 | 10 | Cytoplasm |
| FOXP3 Opal 620 | 6 | Nucleus |
| CD3 Opal 570 | 5 | Cytoplasm |
| CD8 Opal 520 | 0.6 | Cytoplasm |
| PD1 Opal 650 | 4 | Cytoplasm |
| PDL1 Opal 640 | 3 | Cytoplasm |

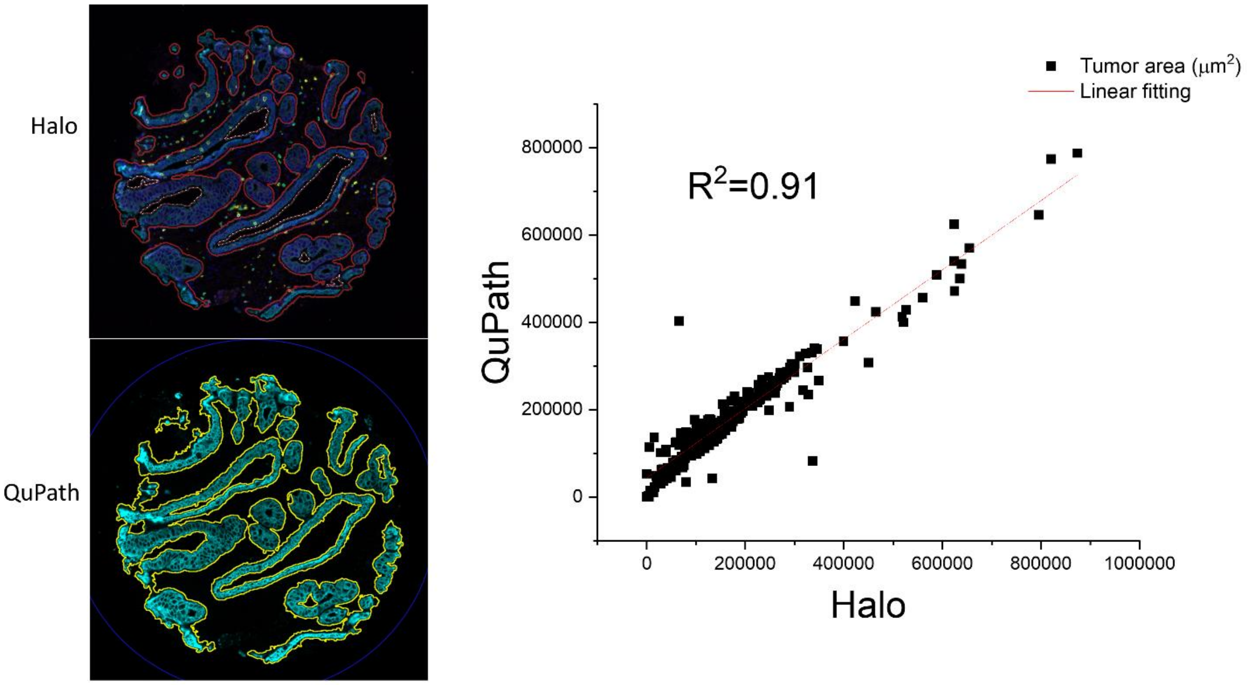


**Supplementary Figure 1.** Tumor segmentation based on pan cytokeratin staining. Both Halo and QuPath software were used for comparison. The tumor area calculated by two programs were analyzed by linear correlation.

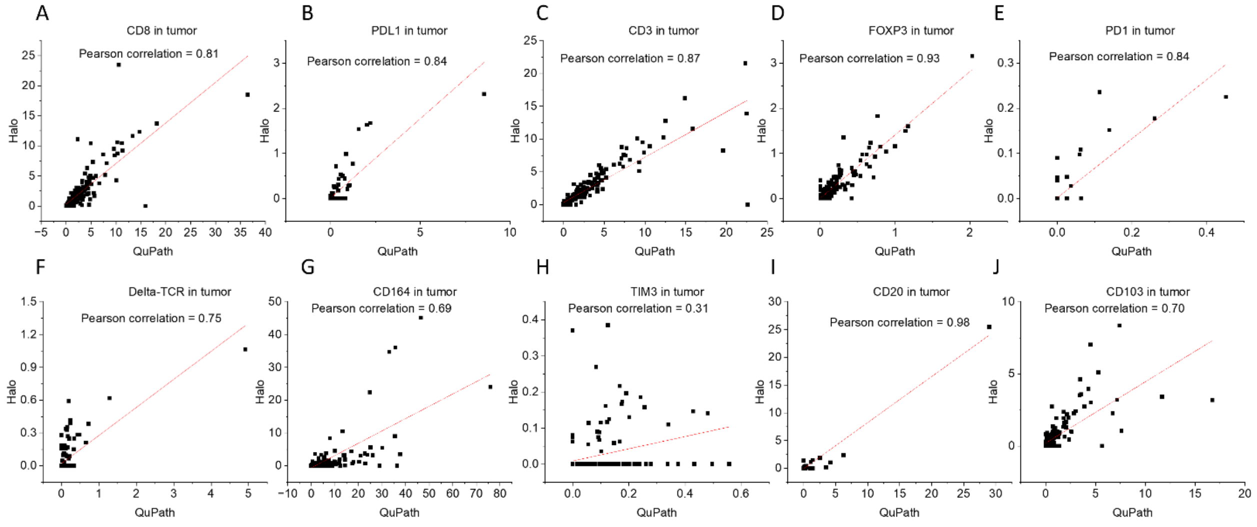


**Supplementary Figure 2.** Correlation analysis of immune cell positive percentage in tumor region

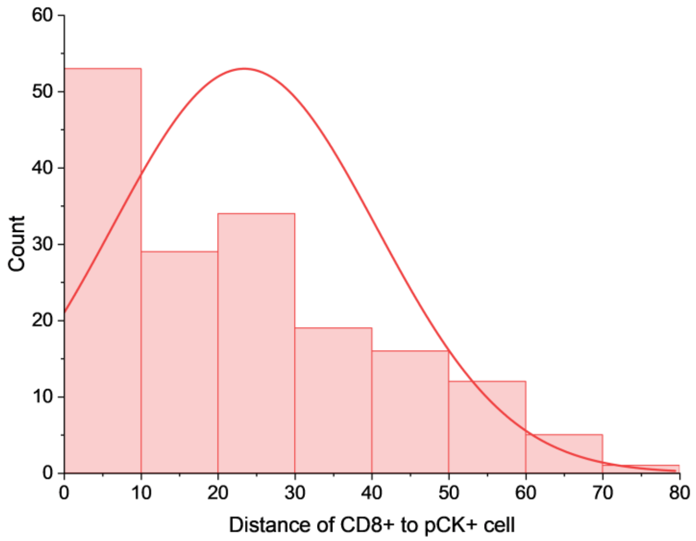


**Supplementary Figure 3.** Histogram of CD8+ cells to nearest pCK+ cells in an example core

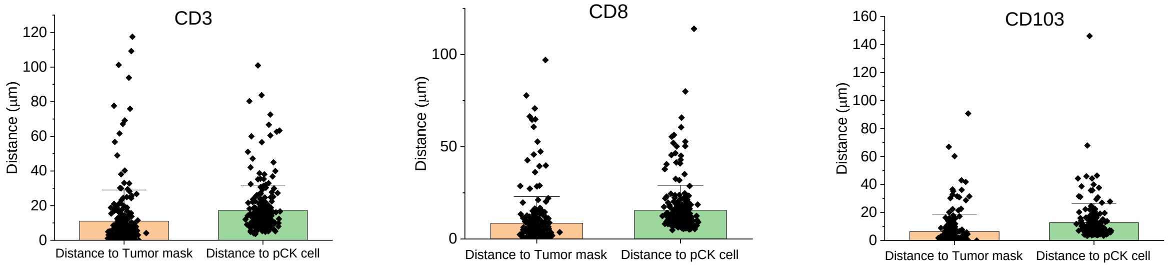


**Supplementary Figure 4.** Comparison of cell to tumor mask distance and cell to cell distance in QuPath.

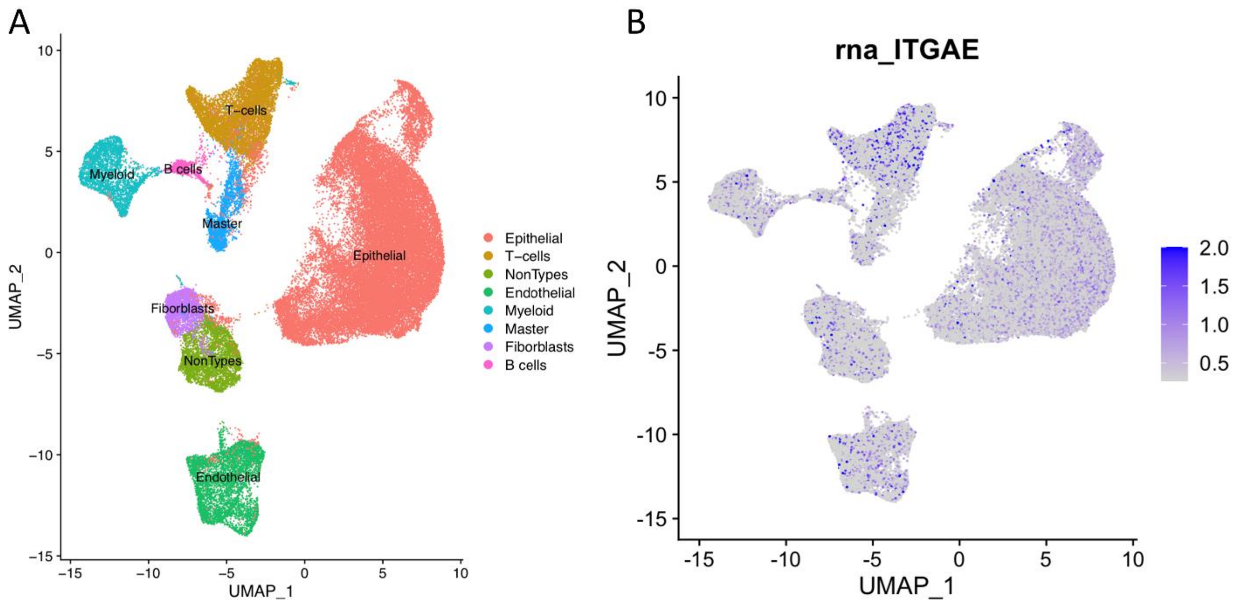


**Supplementary Figure 5.** Single-cell RNA sequencing analysis of distribution of CD103 positive cells in different cell types. A, visualization of single-cell RNA data using Uniform Manifold Approximation and Projection (UMAP), with clustering performed based on well-known gene markers to identify distinct cell types. B, the distribution of CD103-positive cells within each cluster, providing insights into the cellular composition and heterogeneity of the prostate cancer microenvironment, particularly regarding the presence and localization of CD103-expressing cells across different cell populations.
